# Precision and efficacy of RNA-guided DNA integration in high-expressing muscle loci

**DOI:** 10.1101/2024.03.18.582796

**Authors:** Made Harumi Padmaswari, Gabrielle Bulliard, Shilpi Agrawal, Mary Jia, Christopher Nelson

## Abstract

Gene replacement therapies primarily rely on adeno-associated viral (AAV) vectors for transgene expression. However, episomal expression can decline over time due to vector loss or epigenetic silencing. CRISPR-based integration methods offer promise for long-term transgene insertion. While the development of transgene integration methods has made substantial progress, identifying optimal insertion loci remains challenging. Skeletal muscle is a promising tissue for gene replacement owing to low invasiveness of intramuscular injections, relative proportion of body mass, the multinucleated nature of muscle, and the potential for reduced adverse effects. Leveraging endogenous promoters in skeletal muscle, we evaluated two high-expressing loci using homology-independent targeted integration (HITI) to integrate reporter or therapeutic genes in mouse myoblasts. We hijacked the muscle creatine kinase (*Ckm*) and myoglobin (*Mb*) promoters by co-delivering CRISPR-Cas9 and a donor plasmid with promoterless constructs encoding green fluorescent protein (GFP) or human Factor IX (hFIX). Additionally, we deeply profiled our genome and transcriptome outcomes from targeted integration and evaluated the safety of the proposed sites. This study introduces a proof-of-concept technology for achieving high-level therapeutic gene expression in skeletal muscle, with potential applications in targeted integration-based medicine and synthetic biology.

## Introduction

The current wave of therapies transforming genetic medicines rely on gene replacement (*1*). Efficient transgene expression is predominantly achieved through the use of adeno-associated viral (AAV) vectors to express a therapeutic gene under an exogenous promoter (*2*). While multiple studies have reported the sustainability of AAV gene therapy for years after a single administration, others have noted a decline in transgene expression, with some even observing a return to baseline levels (*3–6*). With the majority of transgene expression sustained episomally, research indicates that epigenetic silencing of episomal DNA and interactions between the AAV capsid and host factors significantly contribute to the reduction in expression (*7*). In line with these findings, recent studies in dogs and non-human primates revealed initially high but short-lived expression from episomal genomes, while long-term expression of the transgene was attributed to clonal expansion of cells containing integrated vectors (*8, 9*). However, transgene integration in AAV gene replacement happens in a semi-random pattern throughout the genome (*10*). Vector integration in unpredictable positions in the genome might result in unforeseeable expression and interactions with the host genome, though no adverse events related to vector integration have been detected in humans (*11, 12*). Thus, these limitations highlight the need to transition from random integration with viral vectors to targeted, site-specific methods like CRISPR-Cas technology which can overcome the problem of insertion of correct genes into random genomic sites.

CRISPR-based integration methods have evolved significantly, from using the endogenous repair pathway, non-homologous end joining (NHEJ) or homology-directed repair (HDR), to engineered transposable elements (*13–15*). These gene editing technologies emerged as a potential avenue for long-term therapeutic transgene insertion. Despite considerable advancements in targeted gene integration methods, identifying the locus to insert replacement genes for optimal safety and efficiency remains challenging (*16*). While many gene editing approaches to target the disease locus itself, some studies indicate that the genomic structure of mutated genes may not favor entire gene replacement. Further, some exogenous promoters, such as cytomegalovirus (CMV) and human elongation factor-1 alpha (EF1alpha), are susceptible to promoter silencing (*17, 18*). Therefore, exploring loci with high transcriptional activity emerges as an intriguing alternative.

The current strategy of “hijacking” the endogenous promoter of high transcriptional activity genes or transgene overexpression remains largely organ-specific (*19–21*). While the results are promising, the safety and versatility of this approach have yet to undergo thorough investigation. Ensuring the safety and versatility of these sites is essential for achieving stable expression of integrated transgenes without adversely affecting the host cell. Empirical studies have identified safe-harbor sites that support long-term transgene expression. There are three well-established sites: *AAVS1, CCR5*, and *hROSA26*, and recently explored sites such as *Rogi1, Rogi2*, or *SHS231* (*22–24*). However, the expression from these sites in limited or tissue-specific and requires the inserted vector to include an exogenous promoter.

In this study, we identified new potential safe-harbor sites in skeletal muscle that offer secure and stable integration, facilitating gene replacement. Choosing skeletal muscle as a gene therapy integration site can reduce procedure invasiveness and complexity. Its unique syncytial nature is less likely to create negative effects *in vivo*, with minimal local or systemic adverse effects associated with intramuscular injection were reported in a systematic review for AAV gene therapies (*2*). Owing to its local delivery, the exposure to circulating neutralizing antibodies, such as AAV pre-existing antibodies against the viral capsid is minimized, ensuring efficient tissue transduction (*25, 26*). Skeletal muscle’s well-vascularized nature allows secreted proteins to enter circulation and intramuscular administration of AAV vector that lead to the secretion of functional proteins has been previously demonstrated (*27–29*).

In this study, we devised a strategy to leverage endogenous promoters in muscle cells for expressing either the reporter gene or a therapeutic gene. We adapted the currently established safe-harbor characterization to evaluate two high-expressing skeletal-muscle loci. We applied homology-independent targeted integration (HITI) to mediate transgene integration (*30*), introducing a reporter gene or therapeutic gene (human factor 9, *hF9)* to express human factor 9 protein (hFIX) from the identified loci. By delivering a promoterless transgene, we were able to detect both protein expression in the reporter gene and hFIX. We deeply profiled genome and transcriptome outcomes after gene-editing outcomes using high-throughput sequencing, unidirectional sequencing, and nanopore long-read sequencing. Overall, the data support RNA-guided DNA-integration strategies as effective therapies for restoring desired gene expression in muscle and can be extended to synthetic biology applications.

## Results

### Selected muscle-specific integration sites have high expression and a predicted favorable safety profile

The selection of integration sites involves combining established criteria for potential safe-harbor sites with insights from prior *in vivo* studies that employed the highly expressing locus, albumin (*ALB*), to express therapeutic genes in the liver (*20, 22*). We have tailored this strategy to target integration sites within skeletal muscle. To pinpoint suitable sites, we referred to a comprehensive analysis of public transcriptomics studies on skeletal muscle in humans and mice by Abdelmoez, A. et al (*31*). Through their thorough examination of the compilation and the application of rigorous quality control, normalization, and annotation using official human gene names, we identified the top 10 genes specifically expressed in skeletal muscle and demonstrated high expression levels in both humans and mice (**Table 1**). The selection of these genes is rooted in the goal of ensuring robust and consistent muscle-specific promoter expression in different models of organisms. Next, we evaluated the essentiality of each gene and the safety implications associated with hijacking its endogenous promoter.

**Table 1.** The table compares highly expressed genes in skeletal muscle tissues between mice and humans. The data were extracted from Abdelmoez, A., et al. (2019) (*31*), who compiled skeletal muscle tissue gene expressions from the Gene Expression Omnibus public database. RPKM: Reads Per Kilobase per Million mapped reads, SE: standard error.

|  |  | Human |  | Mouse |  |
| --- | --- | --- | --- | --- | --- |
|  | Gene | RPKM | SE | RPKM | SE |
| 1 | *ACTA1* | 7.03 | 0.27 | 7.14 | 0.14 |
| 2 | *CKM* | 6.91 | 0.24 | 5.65 | 0.60 |
| 3 | *MB* | 6.63 | 0.46 | 6.14 | 0.49 |
| 4 | *MYH7* | 6.72 | 0.33 | 2.26 | 2.80 |
| 5 | *MYL2* | 6.87 | 0.26 | 3.35 | 2.52 |
| 6 | *DES* | 6.11 | 0.44 | 3.35 | 1.68 |
| 7 | *TNNC2* | 6.28 | 0.54 | 6.93 | 0.46 |
| 8 | *TNNI* | 6.31 | 0.43 | 1.93 | 2.73 |
| 9 | *MYL3* | 5.79 | 0.24 | 3.46 | 2.23 |
| 10 | *TTN* | 6.11 | 0.70 | 0.63 | 4.52 |

Skeletal actin (*ACTA1*) is on top of the list as the highest expressed gene in skeletal muscle. However, skeletal actin is the main actin isoform in skeletal muscle and it plays an essential role, alongside myosin, in facilitating muscle contraction(*32*). Further, *ACTA1* disruption in newborn mice causes early demise (*33*). In light of this, we propose creatine kinase (*CKM*) and myoglobin (*MB*) as genomic integration sites for skeletal muscle. While our approach does not involve knocking out the *Ckm* or *Mb* gene, these selections are based on the viability and state of skeletal muscle after a knock-out experiment. Mice lacking *Ckm* are viable and exhibit no changes in absolute muscle force; but, they show an inability to engage in burst activity (*34, 35*). Mice without myoglobin are also viable with no distinct phenotype apart from depigmentation of the cardiac muscle and cardiac adaptations regarding oxygen delivery (*36, 37*).

To identify if these sites could serve as potential genomic safe-harbor sites, we assessed the suitability of creatine kinase and myoglobin as safe-harbor sites in comparison to currently known safe-harbor sites. Scores were assigned based on previously established and widely accepted criteria (**Figure 1**; **Table 2**)(*22*). The table indicates that our proposed sites did not meet all the criteria. However, it is worth noting that even the most commonly used safe-harbor sites fail to achieve a 100% clearance based on these criteria. Interestingly, *MB* shows better results in a free oncogene region criterion compared to *CKM* and other sites.

**Figure 1.**
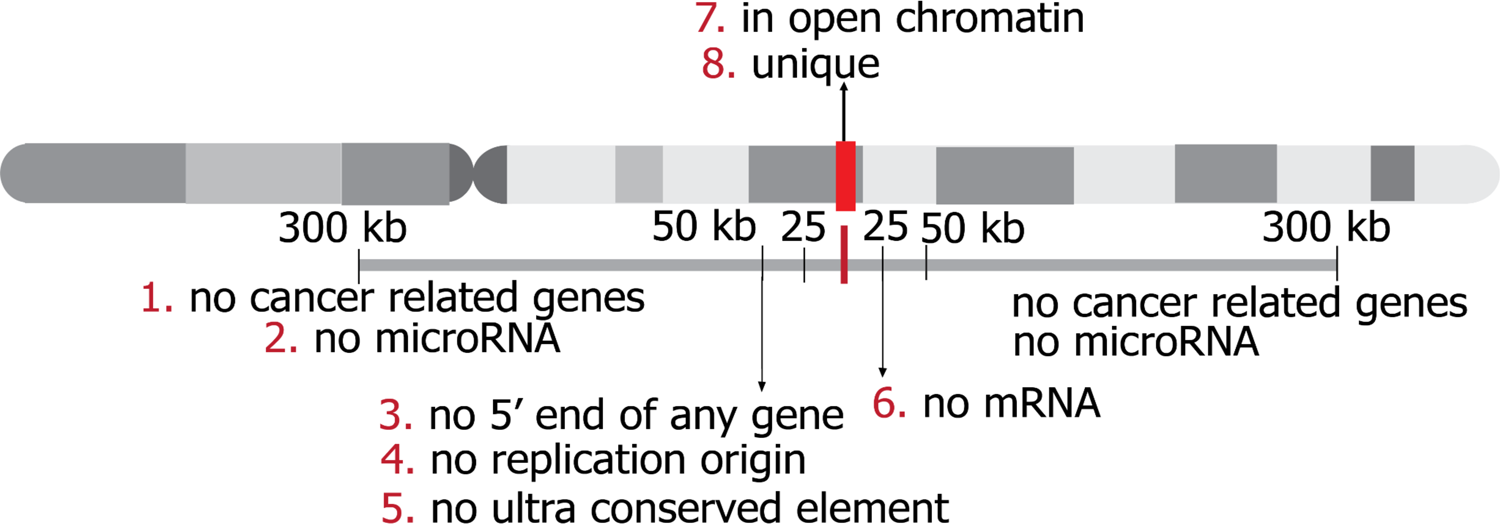
Schematic representation of ideal safe-harbor sites. The red bar in the middle represents the optimal location of safe-harbor sites. The site should be unique and located in open chromatin (*68, 69*). Within 300 kb, the site should be free from any cancer-related genes, miRNAs, or other functional small RNAs (*70*). Within 50 kb, the site should be free from the 5’ end of a gene, replication origin, and ultra-conserved element (*71*–*73*). Additionally, the site should be in a region of low transcriptional activity or have no mRNA within 25 kb.

**Table 2.**
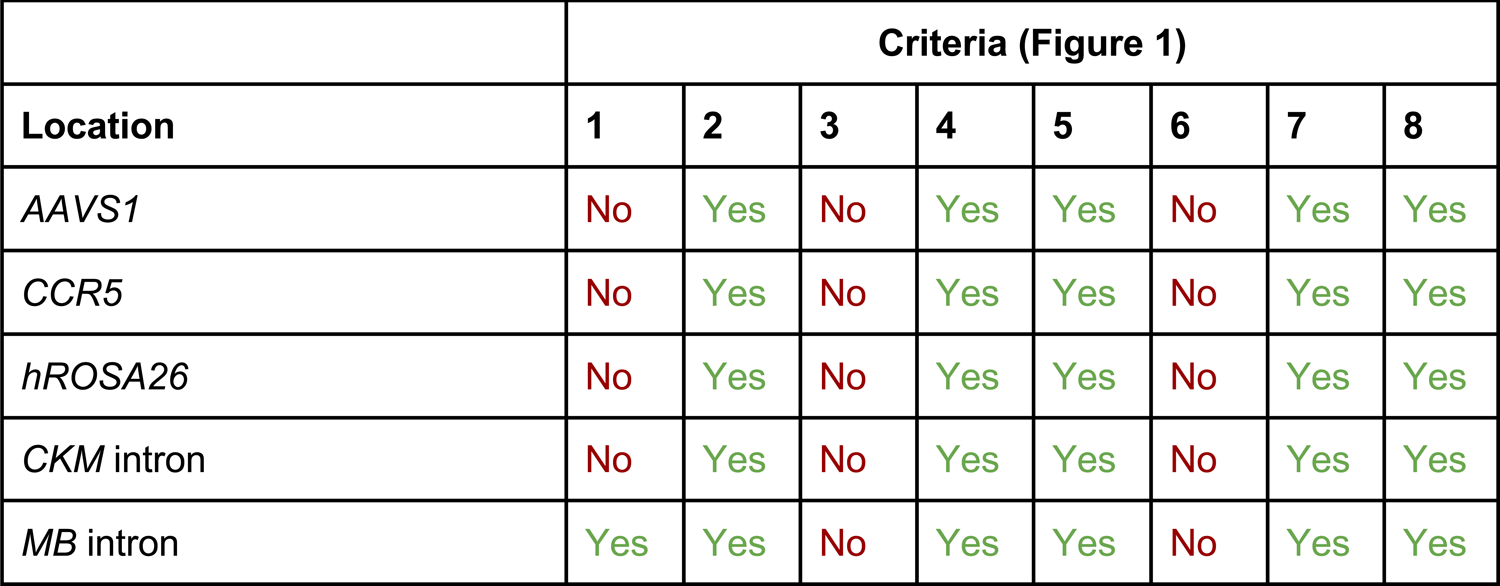
The table displays the assessment of currently widely used safe-harbor sites (*AAVS1, CCR5*, and *hROSA26*) and proposed safe-harbor sites. The assessment was performed based on the ideal criteria of safe-harbor sites. Each criterion was evaluated in the UCSC genome browser track following recommendations from Pellenz, et al.(*22*).

|  | Criteria (Figure 1) |  |  |  |  |  |  |  |
| --- | --- | --- | --- | --- | --- | --- | --- | --- |
| Location | 1 | 2 | 3 | 4 | 5 | 6 | 7 | 8 |
| *AAVS1* | No | Yes | No | Yes | Yes | No | Yes | Yes |
| *CCR5* | No | Yes | No | Yes | Yes | No | Yes | Yes |
| *hROSA26* | No | Yes | No | Yes | Yes | No | Yes | Yes |
| *CKM* intron | No | Yes | No | Yes | Yes | No | Yes | Yes |
| *MB* intron | Yes | Yes | No | Yes | Yes | No | Yes | Yes |

### Targeted integration into *Ckm* and *Mb* lead to expression of a promoterless reporter

To experimentally assess transgene expression from proposed sites (intron 1 of *CKM* or *MB*), we performed a targeted integration approach to knock in a gene construct encoding a promoterless green fluorescence reporter protein (GFP) into mouse myoablsts (*Ckm* or *Mb*). We identified the knock-in site within the intron preceding the coding sequence of the gene, ensuring that the non-edited or inaccurately edited copy maintains functional gene expression via mRNA splicing. This approach also provides flexibility in transgene selection, enabling the incorporation of specific features, such as a signaling peptide or a transgene that does not require a signaling peptide.

To integrate GFP, we used a CRISPR/Cas9-based genome editing strategy that uses the HITI method capitalizing the NHEJ pathway that is accessible in muscle cells (**Figure 2A**). This approach designs Cas9 target sites in the donor DNA as reverse complements of the genomic target site to facilitate the re-cleavage of reverse-integration products, providing a means to dictate the specific directionality of the knock-in. We applied this strategy to C2C12 immortalized mouse myoblast cell lines as it is the most commonly used cellular model to study murine skeletal muscle *in vitro* (*31, 38*).

**Figure 2.**
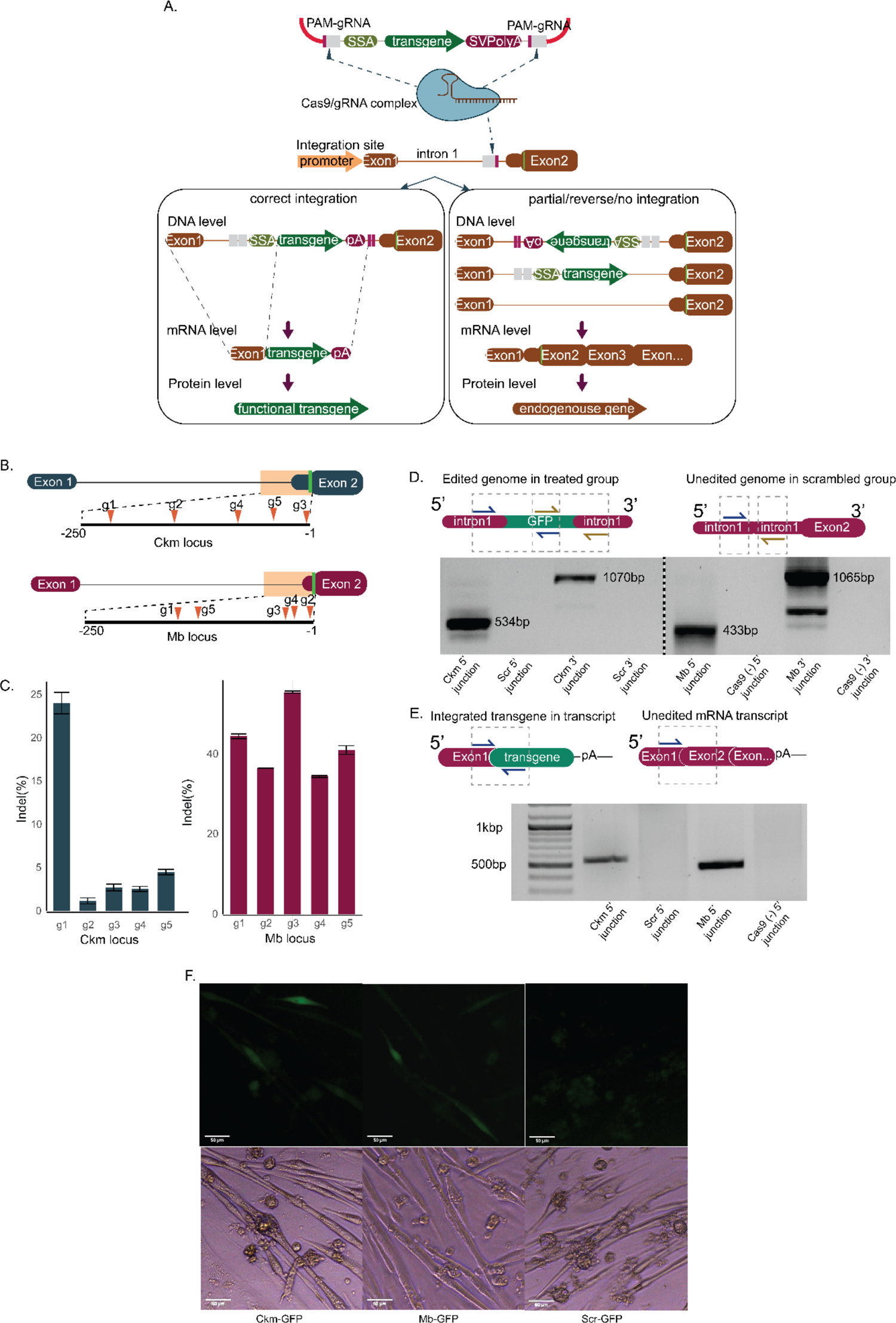
Targeted Integration of a promoterless GFP at Mb and CKm leads to GFP expression in myotubes. **2A.** (Top) Overview of transgene integration to hijack endogenous *Ckm* or *Mb* promoters. The HITI-based gene editing system is delivered via plasmid transfection, which encodes the CMV-driven *S. pyogenes* Cas9 protein, U6 promoter-gRNA expression cassette, and a donor plasmid containing a transgene fragment flanked by Cas9 target sites in the reverse orientation relative to genomic DNA. (Bottom) The schematic mechanism at the molecular level shows that by integrating the transgene in the intronic region, only correct integration is expected to produce the protein, while partial or reverse integration is expected to not affect endogenous genes. The figure is not drawn to scale. **2B.** The gRNA target site map in *Ckm* and *Mb* locus. Five gRNA target sites were selected within 250 bp in the splice acceptor and 5’ UTR region of the gene. **2C.** The quantification of gRNA efficiency in C2C12 cell line with short-read next-generation sequencing (NGS) based on indel percentage. The error bars were computed as the standard deviation obtained through bootstrap resampling technique. **2D.** Validation of correct GFP integration in genomic DNA is shown in both 5’ and 3’ integration regions in the treated group. The schematic figure shows the forward primer in the GFP transgene overlaps with the reverse primer (Primer list). **2E.** Validation of correct mRNA splicing from GFP integration by cDNA PCR is shown in both loci. At the transcriptome level, only 5’ integration was assessed. **2F.** Fluorescent microscopy images assessing GFP expression in edited myotubes at 10 days after editing are shown; the scale bar is 50 µm.

As a proof of concept, we have identified five guide RNA (gRNA) sites within intron 1 of each gene, positioned distal to the enhancer and proximal to exon 2, specifically within the last 250 base pairs leading up to the noncoding portion of exon 2 (**Figure 2B**) based on CRISPOR web tool and screened their cutting efficiency in NIH3T3 cell line (*39*) (**Figure S1**). Additionally, we conducted PCR-enriched next-generation sequencing (NGS) using Iseq short-read sequencing to quantify the efficiency of gRNA in the C2C12 cell line. Within the *Ckm* gene, g1 exhibited the highest efficiency, making it the selected gRNA for HITI-mediated GFP integration experiments. In the *Mb* gene, although g3 demonstrated slightly highest cutting efficiency, it targeted the non-coding exon sequence. Therefore, we opted for g1 as the gRNA for the downstream integration experiment (**Figure 2B and 2C**).

We used Lipofectamine 3000 to co-transfect C2C12 muscle myoblasts with a ratio of 2:2:1 for SpCas9, guide RNA, and site-specific promoterless insert plasmids, respectively. Three days after transfection, we performed a genotyping PCR to confirm the integration. PCR primers were used spanning the intron of *Ckm* or *Mb* to the 5’ junction of the GFP insert, and from the 3’ junction of the GFP insert to the intron of *Ckm* or *Mb*. We detected bands at the expected length of chimeric *Ckm-GFP or Mb-*GFP integration in both 5’ and 3’ end junctions at the genomic DNA and cDNA levels, but no bands were observed in the control groups (**Figures 2D** and **2E**). In myotubes, after differentiation, we observed GFP expression in the CRISPR-treated group in both loci, while the scrambled group did not exhibit GFP expression. This suggests that the GFP expression was driven by the *Ckm* or *Mb* promoter (**Figure 2F**).

### Targeted integration of a promoterless human F9 gene at *Ckm* and *Mb* leads to sustained expression of *hF9*

Next, we sought to investigate whether this strategy applies to therapeutic genes. The GFP transgene in the HITI insert plasmid was substituted with a transgene encoding human factor IX (*hF9*) (**Figure 3A**). The selection of *hF9* as a proof of concept for therapeutic gene integration was motivated by previous reports indicating that even with <1% of targeted integration events of *hF9* under the albumin promoter, it proved adequate to attain 5–20% of FIX levels, effectively correcting bleeding in hemophilia B mice (*21*). In this approach, we utilized the complete coding sequence of *hF9*, given the absence of signaling peptides in the *Ckm* or *Mb* gene. Following a method similar to GFP transfection, we co-delivered a site-specific *hF9* HITI insert plasmid with SpCas9 and *Ckm* or *Mb* gRNA into C2C12 myoblasts using Lipofectamine 3000. Post-transfection, as observed in GFP integration, we verified *hF9* integration exclusively within the CRISPR-mediated HITI group at both genomic DNA and cDNA levels in both sites (**Figure 3B** and **3C**).

**Figure 3.**
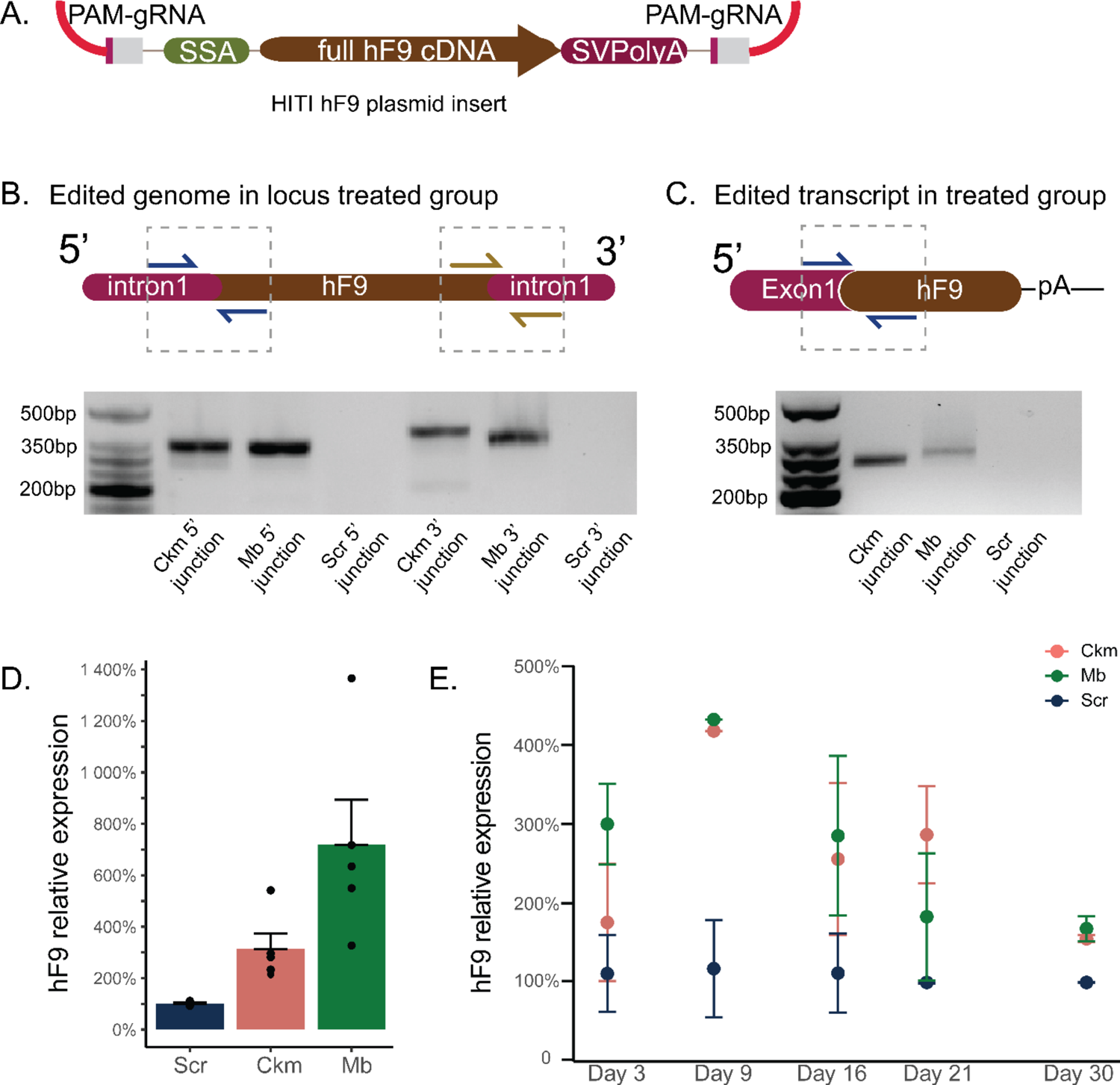
Targeted integration of a promoterless hF9 gene leads to sustained expression 3A. Schematic of the structure of HITI insert plasmid for human F9 expression. Full-length cDNA region from exon 1 to exon 8 is cloned to replace GFP transgene in the previous construct. **3B and C.** Genotyping of the F9 integration in C2C12 cell line using primers spanning the junction between the integration site and the transgene in genomic DNA and cDNA. **3D.** Relative *hF9* expression following plasmid transfection. Five biological independent samples, mean ± s.e.m. All samples were processed at 10 days post-transfection including 7 days of myotubes differentiation. **3E.** Time course cell culture to assess the expression of *hF9* in treated C2C12 for 30 days without selection compared to the scrambled group. Two independents biological replicates, means ± sem.

To evaluate changes in gene expression between groups, we conducted qRT-PCR using FAM probe primers spanning from exon 6 to exon 7 of *hF9*. The *Ppia* gene served as a housekeeping gene for normalizing expression levels. This selection was based on our data that showed the *Ppia* gene has the most consistent expression in both myoblasts and myotubes. In contrast, *Gapdh* has variable expression levels between the myoblast and myotube stages. Furthermore, we evaluated four different reference genes from a previous study to identify the most consistent expression in both myoblast and myotubes (*40*)(**Table S1**).

During the myotube stage, the relative expression of *hF9* was found to be elevated in the CRISPR-mediated HITI group compared to the scrambled group, reaching up to a 3-fold increase in *Ckm*-mediated expression and up to a 10-fold increase in *Mb*-mediated expression (**Figure 3D**). Additionally, to assess if this increased mRNA expression translates to hFIX protein expression, we performed an hFIX sandwich ELISA assay on cell media after myotube differentiation. We observed an increased level of protein expression in both *Ckm* and *Mb*-treated cell serum relative to a scrambled control (**Figure S2**). Furthermore, to determine whether identified sites supporting long-term stable *hF9* expression, we maintained the myoblast culture for 30 days. Despite our efforts to incorporate a reporter gene marker in Cas9 for cell sorting, we encountered difficulties in maintaining the sorted cultured cells. This obstacle has also been noted in previously published literature, which indicates that myoblasts tend to lose their differentiation potential after single-cell cloning, and those that are successfully edited often fail to survive the stress of sorting (*41*). Since generating clonal populations was not feasible, we opted to gather informative data from bulk populations by routinely passaging the cells when they reached 60-70% confluency, before spontaneously differentiating to myotubes.

Following our 30-day culture, we detected integration in DNA level (**Figure S3**). In the time-course analysis, *hF9* gene expression was assessed at various intervals starting from 3 days post-transfection, with cell passaging every two to three days, followed by differentiation from day 23 to 30. Despite the continuous passaging of cells, this longitudinal study demonstrated generally consistent expression of *hF9* throughout the observation period. The variance in fold change observed between fully differentiated myotubes in **Figure 3D** and **Figure 3E** could be attributed to the continuous passaging process, which involved regular splitting of the cells.

### Multi-pronged targeted next-generation sequencing and long-read sequencing show relative precision of HITI-based integration

While we have confirmed the *Ckm* and *Mb* endogenous promoter can express the promoterless protein, the precision of editing was unknown. Imprecise integration could negatively impact protein expression. We evaluated factors affecting protein expression such as the precision and the efficiency of integration. We performed PCR-enriched short-read sequencing for fragments spanning the genomic insertion site or chimeric transcript to evaluate if the integrated chimeric sequence is as precise based on the mechanism of HITI or if unintentional on-target events could be detected. Genomic DNA-level analysis revealed levels of precise integration ranging from 15% to 35% in 5’ and 3’ integration junctions in both loci (**Figure 4A** and **4B**). The modified sequence predominantly resulted from NHEJ repair at gRNA cut sites with preferential removal of sequence on the inserted sequence (**Figure 4A-B**). Although genomic DNA-level precision was not optimal, at the mRNA level, we saw a higher average of precision integration around 80% of precise integration (**Figure 4C**) with some modifications on the junction site of exon 1 *Ckm* and start codon GFP (**Figure 4D**).

**Figure 4.**
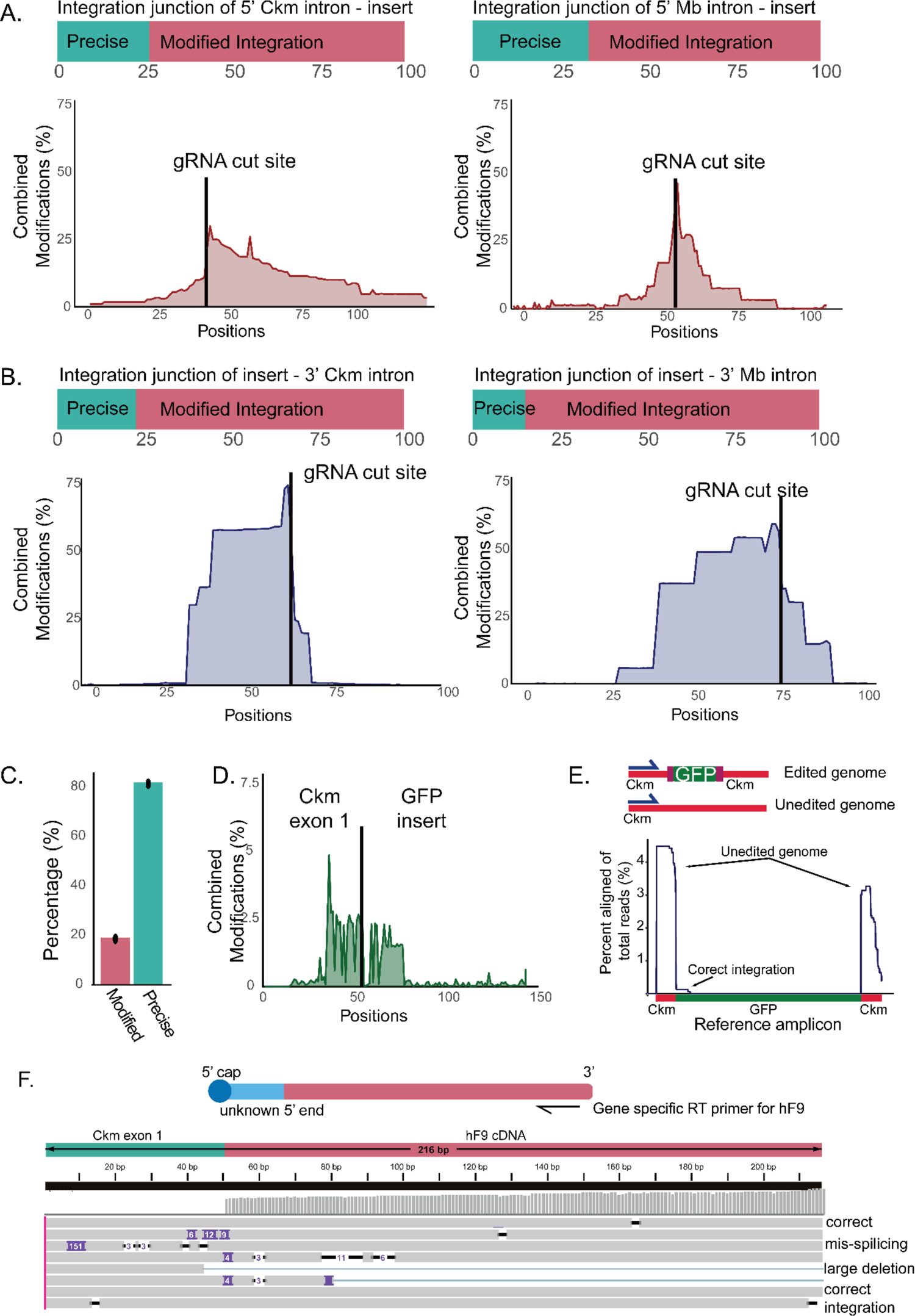
Deep sequencing reveals precise and imprecise outcomes of targeted integration **4A and 4B.** Deep sequencing shows a varied range of precise integration in 5’ and 3’ junctions. A similar pattern was observed in both 5’ and 3’ junctions in both loci, vector chewback that predominantly happened on the insert side. **4C.** Deep sequencing at the mRNA level showed a higher percentage of precise integration in three biological replicates in the treated *Ckm* locus. **4D.** Modification frequency trace from the fusion of *Ckm* and GFP transgene is shown. **4E.** (Top) Schematic Tn5 tagmentation-based sequencing to quantify integration efficiency. (Bottom)The graph shows the percentage of reads aligned to the *Ckm* gene and reference amplicon. **4F.** Characterization of the fusion of *Ckm* and *hF9* cDNA using 5’RACE with GSP reverse transcriptase primer is shown.

Precision integration data provide a selective perspective on how effectively this method incorporates the transgene insert. However, selection of primers and PCR will bias the result. Moreover, this method does not quantify the proportion of integrated sequences due to the PCR-enrichment method that selectively amplifies the integrated sequences. To quantify the integration efficiency, we conducted a Uni-Directional Targeted Sequencing methodology (UDiTaS) that is based on Tn5 transposase-assisted tagmentation short-read sequencing (*42*). This method incorporates a unique molecular identifier (UMI) to remove PCR amplification bias. We quantified the number of sequences integrated with our transgene by tagging primers specific to the *Ckm* intron before the gRNA cut site (**Figure 4E**). Reading from this site will capture unedited, precisly edited, and unintentionally modified alleles. In **Figure 4E**, we deduplicated the UMI and aligned the resulting bam files to the expected fusion of *Ckm* intron-GFP integration. After implementing filters (details in Method) to verify that the plotted reads were authentic and not sequencing artifacts, the analysis revealed that correct GFP integration represented 2.8% of the total aligned reads. This percentage was calculated by dividing the proportion of edited reads by the proportion of unedited reads that aligned with the reference amplicon from the total reads (4%). Separately, a primer specific to the GFP insert genome was used in conjunction with the same transposon-specific primer to map genome-wide GFP episome integration into the mouse genome. Following similar pipeline with UdiTaS analysis, we did not detect GFP integration in other locations in the genome (**Figure S4**).

The cDNA selective PCR-enriched sequencing method revealed a high percentage of precise integration. However, it may overlook large structural variants. Therefore, to get a more comprehensive picture of the precision, we aimed to characterize the 5’ integration junction of the fusion between *Ckm* and the transgene at the transcriptome level. This approach is motivated by the expectation that intron rearrangements would be spliced out in mRNA processing. To achieve this, we conducted 5′ rapid amplification of cDNA ends (RACE) using cDNA from Lipofectamine-transfected cells and amplified *Ckm*-*hF9* fusion transcripts using a gene-specific primer (GSP) positioned at the end of the *hF9* region (*42, 43*). Long-read nanopore sequencing confirmed the addition of exon 1 of *Ckm* in the 5’ region of *hF9*. However, we also observed several structural rearrangement events, including significant deletions and insertions (**Figure 4F**). Each line in the figure represents a single alignment. Upon investigation of the 151 bp insertion, we determined that it originated from the splice acceptor region incorporated from the HITI insert plasmid. This suggests the occurrence of mis-splicing events at the transcriptome level.

### RNA sequencing reveals differentially expressed genes after targeted integration

To investigate whether the *hF9* targeted integration into identified sites resulted in alterations in the overall transcriptome profiles, bulk RNA-seq analysis was performed. After seven days of myotube differentiation, samples exhibiting the high *hF9* expression levels in *Ckm* and *Mb* based on qPCR analysis were compared with scrambled-treated cells from the same experimental batch (**Figure 5A**). Paired-end sequencing on the DNBSEQTM Sequencing System from BGI with an average read length of 100 bp was used for sequencing two biological replicates from each treatment. Principal component analysis (PCA) was performed, followed by visualization of each sample in two dimensions using the first two principal components (PCs). PCA revealed biological variation between samples in the same treatment groups. However, we observed transcriptional similarity within the *Ckm*-integrated and scrambled group and transcriptional variations in *Mb*-treated samples compared to the scrambled group (**Figure 5B**).

**Figure 5.**
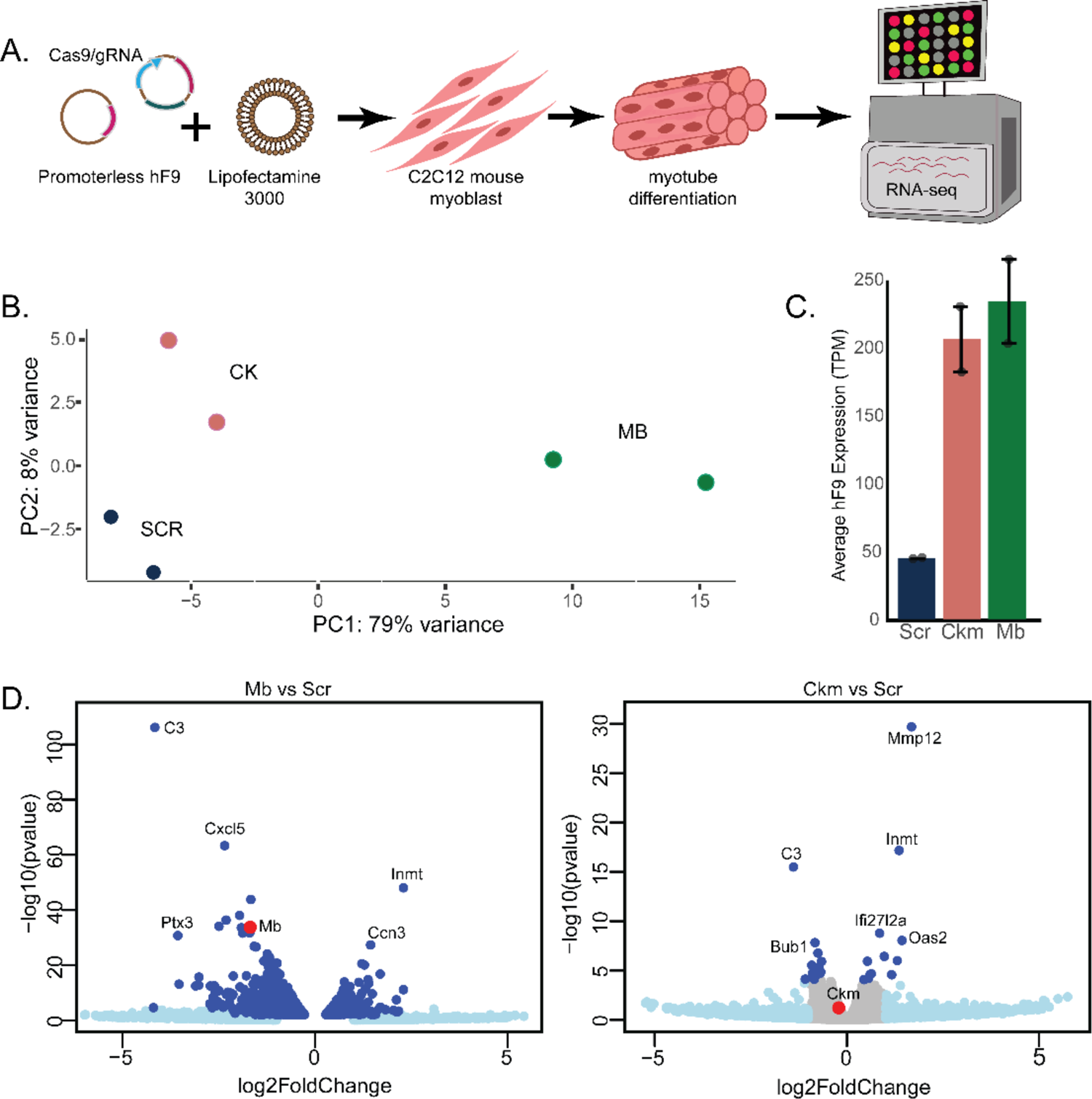
RNA sequencing reveals significant increases in transgene expression and relative precision of integration at *Ckm* over *Mb* **5A.** Pipeline for bulk RNA-seq experiment on *Ckm*- and *Mb*-integrated and scrambled non-integrated C2C12 myotube cells **5B.** Principal component analysis of two biological replicates of C2C12 myotubes with *Ckm, Mb*, or scrambled treatment **5C.** *hF9* transcript expression level measured in TPM is shown, mean ± sem **5D.** Differential expression of genes following *hF9* integration in *Ckm* and *Mb* loci

To assess human *F9* expression in the samples, we normalized the transcript expression level (TPM) and found that *hF9* is highly expressed in both *Ckm* and *Mb* loci compared to the scrambled group (**Figure 5C**). Subsequent differential gene expression analysis was performed to evaluate changes in the transcriptome post-treatment. The analysis revealed several genes that are both downregulated and upregulated in both *Ckm* and *Mb*-treated samples, such as *C3* and *Inmt* genes. Differentially expressed genes were observed to be dispersed across various chromosomes (e.g., *C3* on chromosome 17, *Inmt* on chromosome 6, and *Mmp12* on chromosome 12), rather than being clustered within the targeted chromosome (*Ckm* on chromosome 7; *Mb* on chromosome 15), where more local contacts are expected to occur. Consistent with the PCA results, more genes were differentially expressed in *Mb*-treated samples compared to *Ckm*-treated samples (**Figure 5D**). Additionally, downregulation of the *Mb* gene was observed upon editing.

## Discussion

Our current work aims to address several significant challenges associated with AAV vector-based *in vivo* gene therapy, including long-term efficacy, promoter or transgene silencing, and the safety of transgene integration (*10, 44, 45*). To enhance the sustainability of the treatment, targeted integration into the host genome has emerged as a potential solution. However, the location of where the integration must be strategic since the integrating vector with the constitutive promoter might pose risks. Despite their ability to facilitate robust and consistent transgene expression, such promoters are associated with increased risks, including an elevated likelihood of inactivation, amplified toxicity resulting from transgene overexpression, off-target transgene expression, and the potential for severe immune responses due to inadvertent transgene expression in antigen-presenting cells (*45*).

Utilizing endogenous promoters for transgene expression can circumvent potential silencing mechanisms associated with exogenous promoters. In this study, we identified two highly expressed genes, *Ckm* and *Mb*, in skeletal muscle and exploited the transcriptional output of their endogenous promoters. Our findings demonstrate that these loci can drive the expression of promoterless GFP and enhance RNA expression of promoterless *hF9* in C2C12 cells upon plasmid transfection using lipofection. However, due to the limited efficiency of lipofection in myoblasts, compounded by the generally low efficiency of integration, we confirmed that the efficiency of transgene integration at the genomic level is approximately 3% through the UDiTaS sequencing approach (*46, 47*). Despite this modest level of integration, we observed the expression of promoterless GFP and increased RNA expression of *hF9* compared to a control, representing the episomal vector without Cas9.

Furthermore, it is noteworthy that the expression levels of *Ckm* and *Mb* in C2C12 cells were not as high as reported in skeletal muscle tissue (*31*) (**Table S2**). This disparity may contribute to the lower expression of GFP and hFIX protein observed. Additionally, regarding the hFIX protein results, the rich presence of collagen IV in skeletal muscle may facilitate the local attachment of FIX, limiting its release into circulation (*48*). Previous studies exploring the production of ectopic hFIX protein in muscle have suggested the potential use of hFIX variants harboring mutations such as lysine to alanine at residue 5 (K5A) or valine to lysine at residue 10 (V10K), which exhibit reduced binding to endothelial cells while maintaining normal clotting activity, thereby enabling synthesis of hFIX in skeletal muscle (*49, 50*). Hence, investigating the expression of this variant under a muscle endogenous promoter in muscle cells could potentially result in increased hFIX release into circulation.

The general safety evaluation of the proposed sites indicates that the group treated with *Mb* exhibits alterations in the global transcriptome. The downregulation of the *Mb* gene may be attributed to the higher indel activity of the CRISPR/gRNA complex at that target site compared to the indel activity at the *Ckm* target sites. Despite observing minimal changes in the expression of upregulated and downregulated genes at the *Ckm* locus, which may suggest passing initial safety validation, a more comprehensive analysis is still warranted. *C3* downregulated expression was reported in the acute stage of infection suggesting that *C3* participates in inflammation and slight upregulation of *Inmt* was reported in skeletal muscle under spaceflight conditions (*51, 52*). Further investigation is required to establish a direct correlation between transgene integration and the observed differential expression. However, it is important to note that these *in vitro* analyses may not accurately reflect CRISPR-mediated integration activity *in vivo*. Differences in intracellular nuclease levels and chromatin states between *in vitro* and *in vivo* conditions can potentially impact editing activity at both on- and off-target sites. Further investigations are required to elucidate these differences and to assess the safety and efficacy of CRISPR-mediated integration *in vivo* (*53*).

Our sequencing studies also show patterns of deletions following integration induced by double-strand breaks (DSBs). These sequences demonstrate that indel-induced damage tends to occur toward the insertion site, as evidenced by the selected PCR-enriched integration sequencing. This trend is consistently observed across loci and in both 5’ and 3’ integrations. This insight is valuable for future template design efforts, emphasizing the importance of incorporating padding sequences before the coding sequence of the transgene to mitigate chewback into the desired gene before integration can occur.

Our approach integrating the transgene into intronic regions offers the advantage of splicing, as indels occurring at the intronic level are typically spliced out, resulting in more precise mRNA. Another study has also reported the advantages of intron targeting, which results in the production of nearly error-free mRNA (*54*). However, identifying the strongest or most optimal splice acceptor poses a significant challenge. Despite selecting a robust splice acceptor from a highly conserved gene in skeletal muscle, based on our 5’ RACE results, we observed instances of missplicing due to the failure of the splice acceptor to be properly excised. To date, there is a lack of practical guidelines available to determine the ideal length of splice acceptors or splice donors for a given gene. Thus, design of the splice acceptor allows additional optimization of gene expression.

The two sequencing approaches that we performed, both short and long read, provided us with a general overview of the effects of double-strand breaks (DSB) on integration sites. As targeted integration gains prominence as a viable gene editing therapy approach, we should not overlook the analysis of post-DSB-induced integration events. A more thorough analysis regarding structural rearrangements and vector integration is needed to get a full understanding of these consequences. Another potential avenue involves exploring recent DSB-less or non-DSB approaches, such as ShCAST, TwinPE, PASTE, CAST, HELIX, and PRINT (*55–60*).

In summary, this work presents a proof-of-concept technology for achieving high-level therapeutic gene expression in skeletal muscle. This method holds promise for advancing targeted integration-based medicine or synthetic biology approaches, effectively transforming skeletal muscle into a biofactory capable of producing desired therapeutic agents. While further refinement is necessary to enhance several aspects of the concept, the findings of this research offer a potential avenue towards realizing the promise of gene editing therapy, thereby extending its reach to broader populations.

## Material and Methods

### *In vitro* gRNA screening

gRNA plasmids were constructed by cloning the targeted sequences into the pX330 vector (Addgene #42230) via the BbsI sites under the human U6 promoter. A panel of gRNAs was designed based on high efficiency and predicted minimal off-target effects using the CRISPOR tool from UC Santa Cruz to target intron 1 in each site (*39*). The SpCas9 protein is expressed in a separate plasmid under the CMV promoter. gRNA activity was compared by surveyor assay in NIH3T3 and by next-generation sequencing (NGS) in C2C12 cell lines. NIH3T3 cells were obtained from ATCC (#CRL-1658) and maintained in Dulbecco’s modified Eagle’s medium (DMEM) (Gibco) supplemented with 10% fetal bovine serum (FBS) (Sigma) and 1% penicillin-streptomycin (P/S) (Gibco). C2C12 cell lines were obtained from ATCC (CRL-1772) and cultured in DMEM supplemented with 20% FBS and 1% P/S. Both cell lines were grown in a tissue culture incubator at 37°C and 5% CO2. NIH3T3 cells were transfected with the plasmids encoding SpyCas9 or SauCas9. After a 72-hour incubation period, genomic DNA was isolated using a DNeasy kit (QIAGEN). Indels were identified by PCR of the region of interest (**Tables S3**), followed by incubation with the Surveyor® Nuclease and electrophoresis on TBE agarose gels as previously described. C2C12 transfection was performed using Lipofectamine3000 (Thermo Fisher). In 24-well plate, C2C12 cells were transfected using 250ng pSpCas9 and 250ng pgRNA plasmid. Three days after transfection, cells were harvested and gDNA was extracted as described above. In the second PCR round, full Illumina flowcell adapter sequences along with experiment-specific barcodes were appended to the 5’ and 3’ end of the PCR product, respectively (**Table S3**). After pooling the resultant PCR products, sequencing was performed with 150 bp paired-end reads on an Illumina ISeq instrument (Iseq 100 i1 Reagent v2, 300-cycle). Samples were demultiplexed based on assigned barcode sequences, and the Illumina adapter sequences were trimmed from the reads. The indel analysis was carried out using CRISPResso analysis, following the default parameters (*61*).

### HITI insert plasmid preparation and C2C12 transfection

Plasmid constructs were produced via either gene synthesis or conventional subcloning methodologies. To promote transparency and accessibility, all constructs not previously disclosed, along with their associated sequences, will be submitted to Addgene. Depending on the particular downstream experiments, the donor constructs may include either promoterless GFP or *hF9* coding sequences, both of which are surrounded by the same gene-targeting cutting site, along with splice acceptor and SVpolyA. Exon 1 of *hF9* was ordered from IDT as a gene block, and exon 2-8 was cloned from Addgene #182141. HITI insert plasmids were designed as previously reported(*30*). Cloned plasmids were confirmed with Sanger sequencing, and confirmed plasmids were isolated and purified using Qiagen’s Plasmid Plus Maxi kit following the manufacturer’s protocol.

Cell culture transfections were carried out using Lipofectamine 3000 (ThermoFisher) following the standard protocol. Briefly, 24 hours before transfection, 25,000 C2C12 cells were seeded into 24-well plates to achieve 50-60% confluency the next day. On the day of transfection, two hours before transfection, the growth medium was replaced with a fresh growth medium. Three plasmids: Cas9, gRNA, and HITI insert were first mixed with a ratio of 2:2:1 with the reagent P3000 (ratio of DNA: P3000 of 1:2) in Opti-MEM™ Reduced Serum Medium (Gibco™), and then added to Opti-MEM™ Reduced Serum Medium (Gibco™) containing Lipofectamine®3000. DNA/RNA: reagent complexes were briefly vortexed, kept at room temperature for 15 minutes according to the supplier’s instructions, and added dropwise onto the cell monolayer. Three days post-transfection, when confluency exceeded 90%, myoblasts were differentiated with DMEM supplemented with 2% donor equine serum, 1% Penicillin/Streptomycin, and insulin. Cells were differentiated for 7 days and then harvested. For long-term culture, after transfection, cells were passaged and harvested every two days for 23 days before reaching 80% confluency to prevent spontaneous myotube formation. On day 23, cells were differentiated into myotubes as described above.

### Genomic DNA, RNA, and protein analyses

gDNA extraction from the cell pellet was carried out using a DNAeasy Qiagen kit (#69504) according to the manufacturer’s protocol. RNA extraction was performed using New England Biolab’s Monarch Total RNA Miniprep Kit (#T2010) following the manufacturer’s instructions, including DNAseI treatment. mRNA was then reverse-transcribed using LunaScript RT-supermix (NEB #E3010). Genotyping PCR was then conducted using Q5 polymerase (NEB) for genomic DNA and cDNA. qPCR analysis for *hF9* was executed using Luna® Universal Probe qPCR Master Mix (#M3004) with FAM probe primers (IDT), while the *Ppia* reference gene was assessed using Luna® Universal qPCR Master Mix (#M3003). For primer location, sequences, and combinations, please refer to **Table S3**.

For precision deep sequencing analysis, PCR-based genotyping of *Ckm* and *Mb* sites using primers spanning the 5’ and 3’ junctions of the targeted site and the inserted GFP transgene was performed for 15 cycles. The second round of PCR and the analysis were performed as previously described. In CRISPResso2 analysis, the amplicon sequence was the expected precise integration based on HITI integration (*61*). The percentage of unmodified sequences was tallied as precise integration, while the percentage of modified sequences was categorized as non-precise integration and then mapped to identify the regions where modifications predominantly occurred.

For protein analysis, cell supernatant was collected on days 3 and 7 of differentiation and stored at −80 before analysis. Analysis of *hF9* protein was performed using the Human Coagulation Factor IX Total Antigen ELISA Kit (Innovative Research) following the supplier’s protocol, with 1M HCl used as a stop solution.

### Tn5-mediated tagmentation sequencing

Unloaded Tn5 transposase proteins were acquired from Diagenode (#C01070010-10). Tn5 transposase was then loaded with oligos containing mosaic ends incorporating 13 base-pair unique molecular identifiers (UMIs) and the i5 sequencing adapter according to the manufacturer’s protocol. Tagmentation of 50 ng genomic DNA (gDNA) with 1:8 dilutions of loaded Tn5 was carried out as previously described (*46*). Nested PCRs were then performed using the full i5 adapter and a gene-specific primer positioned in the intron of *Ckm* to enrich the target site and incorporate sample barcodes/the i7 adapter. All PCR was performed with Q5 DNA Polymerase, with an annealing temperature is 60C for 15 seconds and an extension temperature of 68C for 1 minute. Sequencing was conducted on the Illumina iSeq platform using the 150 bp paired-end reagent cartridge following standard protocol.

Sequencing FASTQ files were demultiplexed using the list of barcodes assigned to each sample. Trimmomatic (version 0.33) was then used to trim sequencing adapters and remove low-quality reads (*62*). UMI sequences were annotated using UMI-tools (*63*). Afterward, the annotated FASTQ files were aligned to a reference amplicon using bwa-mem2 (*64*). The reference amplicons were constructed to align with the targeted locus and anticipated edits. SAM files resulting from the alignment were converted to BAM files and then indexed. Deduplication of the indexed BAM reads was performed based on the UMI annotation using UMI-tools. Following deduplication, reads were filtered using seqkit tools to eliminate reads attributed to false priming (reads lacking the 20 bases directly adjacent to the Gene-Specific Primer (GSP) expected sequence were filtered out). Additionally, reads failing to extend adequately beyond the edit site and those falling short of the minimum required length were filtered out. Finally, the deduplicated and filtered reads were analyzed and mapped to quantify the frequency of integration.

### 5’RACE and long read sequencing

The 5’ RACE protocol using the template-switching reverse transcriptase enzyme mix from NEB was executed following the manufacturer’s guidelines. Briefly, 1 µg of RNA was annealed with a gene-specific RT primer containing dNTPs at 70°C for 5 minutes. Next, the mixture was supplemented with template-switching buffer, template-switching oligo, and RT enzyme mix (NEB #M0466), followed by an incubation step at 42°C for 90 minutes and then at 85°C for 5 minutes. Touchdown PCR amplification was carried out using gene-specific reverse primer and TSO-specific primer on 1 µL of the template-switching sample. To assess the enrichment of the amplified fragment in the targeted gene, qPCR was performed on the bead-purified PCR product. Qubit measurement was used to determine the concentration of the purified PCR product. A comparative analysis of qPCR results for cDNA with similar amounts served as a control. A Cq value of <10 was considered indicative of good enrichment for RACE amplification. Enriched samples were then prepared for Oxford Nanopore long-read sequencing library preparation following the manufacturer’s protocol (SQK-NBD112.24), and sequencing was conducted using a MinION flow cell (R10.4.1). Fastq reads were aligned to reference amplicon that has the fusion of *Ckm* exon 1 with the *hF9* coding sequence using minimap2 alignment (*65*), then visualized using Integrative Genome Viewer (IGV) (*66*)

### RNAseq analysis

After qPCR analysis, the best-expressing samples in *Ckm* treated and MB treated, as well as the scrambled samples of C2C12 cells, were further assessed for RNA integrity using Tapestation with High Sensitivity RNA ScreenTape (#5067-5579). RNA samples with RIN > 7 and a 260/280 ratio of around 2 were selected for sequencing. The RNA-sequencing assay was conducted by BGI Genomics (Shenzhen, China). Following RNA sequencing, the FASTQ data underwent quality control using fastqc, and poor-quality reads were trimmed using Trimmomatic (version 0.33) (*62*). High-quality reads were then aligned to mouse and human genomes using HISAT2 (*67*). The number of raw reads associated with each gene was quantified using featureCounts. Next, the raw read data were subjected to principal component analysis and differential expression analysis using DESeq2 in R.

## Acknowledgment

The authors would like all members of the Nelson lab and thank Drs. Nicholas Greene and Kevin Murach for helpful conversations about muscle biology. This work was supported by an NIH/NIBIB R00 #EB023979, ASGCT Career Development Award, University of Arkansas Chancellor’s Innovation Grant, and The Arkansas Bioscience Institute. CEN was supported by the 21^st^ Century Chair in Biomedical Engineering. MHP was supported by the Fulbright Indonesia Research Science and Technology (First) Master’s Degree Program and PhRMA Foundation 2023 Predoctoral Fellowship in Drug Discovery. GNB was supported by the University of Arkansas Honors College Undergraduate Research Grant, MSJ was supported by a State Undergraduate Research Fellowship (SURF), the Bodenhamer Fellowship, and the Goldwater Scholarship.

## Author contributions

MHP and CEN designed experiments and wrote the manuscript. MHP performed experiments, analyzed data, and created figures. MHP, GB, and SA performed molecular cloning and molecular analysis. MHP and MSJ performed short read sequencing.

