## Supplemental Information for "Precision and efficacy of RNA-guided DNA integration in high-expressing muscle loci"

### Supplementary Information

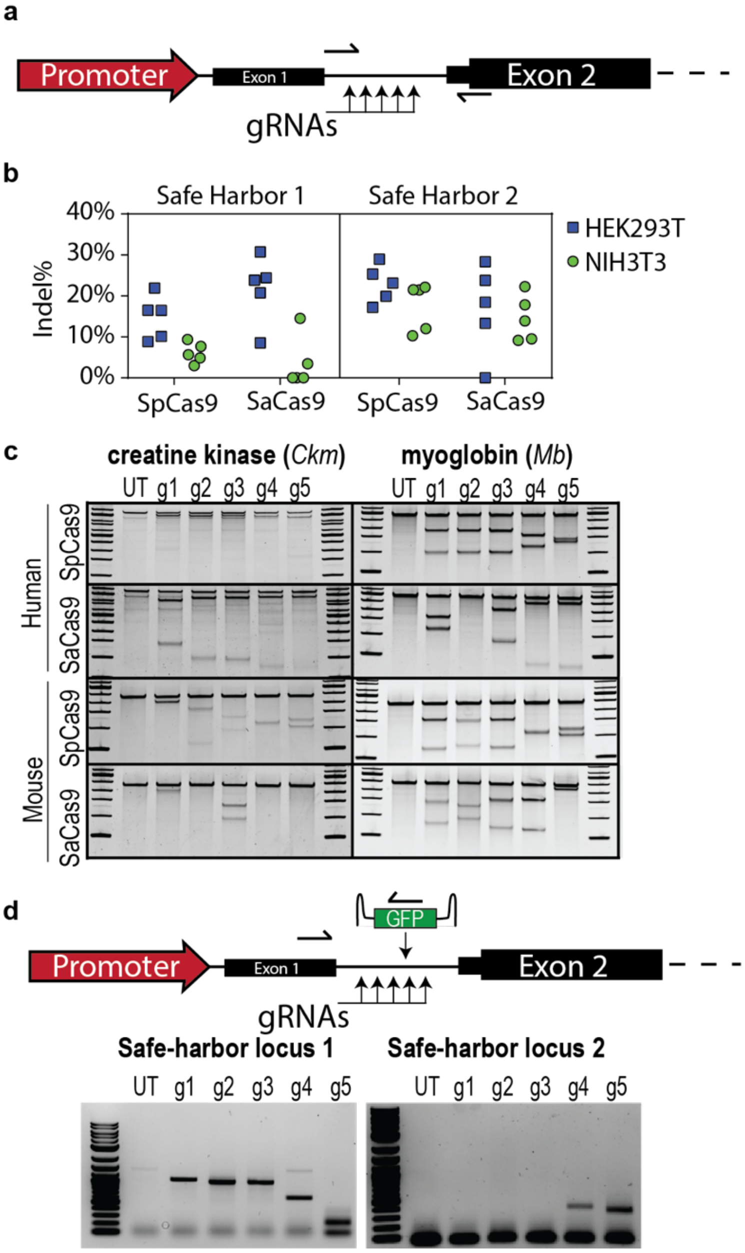


**Figure S1 - T7 surveyor assay of gRNA efficiency in NIH3T3 and HEK293T**. A) A panel of gRNAs targeting creatine kinase in mouse and human were designed targeting intron 1 in each gene. B) SpCas9 and SaCas9 guides were examined for activity by a surveyor assay. C) Gel electrophoresis used of the surveyor assay. D) An AAV encoding GFP was introduced simultaneously to profile targeted integration of the GFP encoding gene.

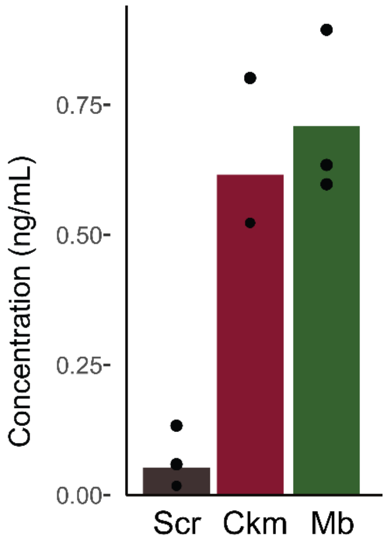


**Figure S2 – Human Factor IX ELISA shows significant increases over scrambled control.** C2C12 cells were transfected with Cas9, *Ckm* or *Mb* gRNA, and donor (promoterless cDNA of *hF9*). Three days post-transfection, cells were differentiated into myotubes. Their media was changed in day 4 and 7 of differentiation, with aliquots taken at day 7. Human Factor IX was present at detectable levels by ELISA in two of three biological replicates in the treated group. hFIX expression was detected below detection level (0.1 ng/mL). Values reflect the mean of two or three independent biological replicates.

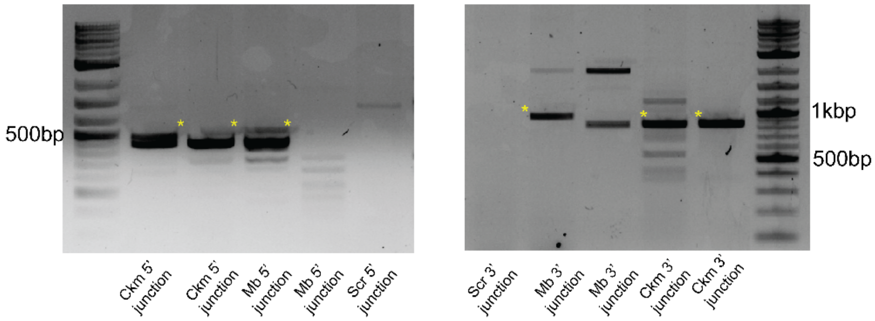


**Figure S3.**
Genotyping of DNA integration in long term culture (day 30) of *Ckm* and *Mb* treated cells integrated with *hF9*. Asterisk represents the expected size of integration.

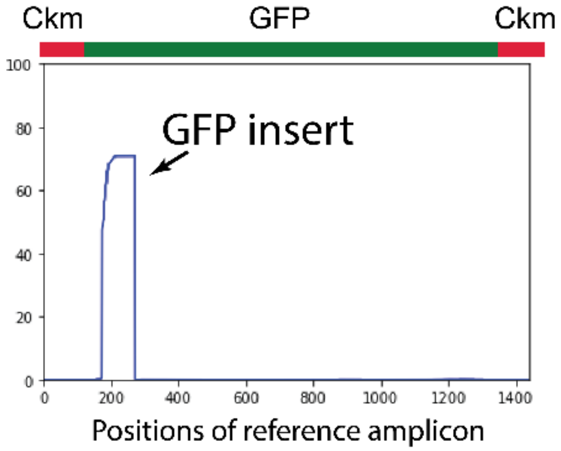


**Figure S4**

A GSP targeting the middle splice acceptor in the GFP insert plasmid was used as an anchor for the UDiTas primer. To assess whether the GFP insert was integrated randomly throughout the genome, fastq reads were aligned to the mouse whole genome using the pipeline described in the Methods section, resulting in no hits when plotted. Furthermore, to determine if the GFP insert was integrated in a different direction than expected, fastq reads were aligned to the reference amplicon. The plot showed that the reads were in the correct direction. However, it is important to note that the depth of sequencing needs to be increased for a more comprehensive analysis.

**Supplementary Tables:**

**Supplementary Table 1. Reference genome comparison Cq value**

| Target | Cq mean | Stage |
|---|---|---|
| *Hprt-1* | 21.73 | Myoblast |
| *Hprt-2* | 21.21 | Myoblast |
| *Actb-1* | 24.81 | Myoblast |
| *Actb-2* | 25.11 | Myoblast |
| *Ppia-1* | 16.89 | Myoblast |
| *Ppia-2* | 16.4 | Myoblast |
| *Gapdh-1* | 16.08 | Myoblast |
| *Gapdh-2* | 16.03 | Myoblast |
| *Hprt-1* | 19.65 | Myotubes |
| *Hprt-2* | 19.5 | Myotubes |
| *Actb-1* | 18.44 | Myotubes |
| *Actb-2* | 18.62 | Myotubes |
| *Ppia-1* | 16.08 | Myotubes |
| *Ppia-2* | 16.01 | Myotubes |
| *Gapdh-1* | 22.48 | Myotubes |
| *Gapdh-2* | 22.38 | Myotubes |

**Supplementary Table 2:**
**RNAseq compilation of mouse and human skeletal muscle (doi: 10.1152/ajpcell.00540.2019)**

| | Human | | Mouse | | C2C12 | |
|---|---|---|---|---|---|---|
| **Gene** | **Amount** | **SD** | **Amount** | **SD** | **Amount** | **SD** |
| *ACTA1* | 7.03 | 0.27 | 7.14 | 0.14 | 6.44 | 1.19 |
| *CKM* | 6.91 | 0.24 | 5.65 | 0.6 | 1.23 | 0.59 |
| *MB* | 6.63 | 0.46 | 6.14 | 0.49 | 3.78 | 1.92 |
| *MYH7* | 6.72 | 0.33 | 2.26 | 2.8 | 1.04 | 1.09 |
| *MYL2* | 6.87 | 0.26 | 3.35 | 2.52 | 1.42 | 2.1 |

**Supplementary Table 3:**
**gRNA list**

| | |
|---|---|
| *Ckm*-1 | AGTCCCCAGCAGGATCACAT |
| *Ckm-2* | GCAAGGCTGAGGTTCACAGG |
| *Ckm-3* | GTGTTGCCGAACGGCATGG |
| *Ckm-4* | ACTTAAGAACTCAGTTGCT |
| *Ckm-5* | GTCTGCTCGCAGGGTCCCAA |
| *Mb*-1 | GCCAGACTAGCATCTGGGA |
| *Mb-2* | CCCATCACTGAGCCCCATGG |
| *Mb-3* | TCCTCTTTAGAAGCCACCA |
| *Mb-4* | CCCCATGGTGGCTTCTAAAG |
| *Mb-5* | GAAGGTATAAAAGCCCTTC |

**Supplementary Table 4 - Primer list**

| | |
|---|---|
| **DNA integration confirmation** | |
| Fwd primer 5' integration *Ckm*-GFP | GAGACCTGTGTAGGAGCCCA |
| Fwd primer 5' integration *Mb*-GFP | CTAGGGGTCAGCAAGATGCC |
| Rev primer 5' integration *Ckm-*/*Mb-*GFP | CCGGACACGCTGAACTTGTG |
| Rev primer 3' integration *Ckm*-GFP | AGCAGACCGGGTTCAGACT |
| Rev primer 3' integration *Mb*-GFP | TCCACCTTCCCCCAGACATTC |
| Fwd primer 3' integration *Ckm-*/*Mb-*GFP | CACAAGTTCAGCGTGTCCG |
| Fwd primer 5' integration *Ckm-hF9* | CAGCCCCAGATCCTGTATTTTTG |
| Fwd primer 5' integration *Mb-hF9* | CCAAAATAGCTGCCCATGTGA |
| Rev primer 5' integration *Ckm-*/*Mb-hF9* | ATCAGCGCAGAAGTCTCAG |
| Rev primer 3' integration *Ckm-hF9* | GGTTCTCCCCCTGTGAACCT |
| Rev primer 3' integration *Mb-hF9* | GCCCCATGGTGGCTTCTAAA |
| Fwd primer 3' integration *Ckm-*/*Mb-hF9* | TCTTAGTCAGGCTGCCCCTC |
| **cDNA integration** | |
| Rev primer cDNA *Ckm-*/*Mb-*GFP | CTCAGGTAGTGGTTGTCGGG |
| Fwd primer cDNA *Ckm*-GFP | ACCTCCACAGCACAGACAGAC |
| Fwd primer cDNA *Mb-*GFP | GTCCCAGGAGAAAGACCCAAT |
| Rev primer cDNA *Ckm-*/*Mb-hF9* | GGACTCACACTGATCTCCATC |
| Fwd primer cDNA *Ckm-hF9* | CCTCCACAGCACAGACAGAC |
| Fwd primer cDNA *Mb-hF9* | CATCCTTGTCCCTGTGGGTG |
| **Iseq primers** | |
| Fwd primer 5' integration *Ckm*-GFP | TCGTCGGCAGCGTCAGATGTGTATAAGAGACAGCAGCCCCAGATCCTGTATTTTTG |

| | |
|---|---|
| Fwd primer 5' integration *Mb*-GFP | TCGTCGGCAGCGTCAGATGTGTATAAGAGACAGCCAAAATAGCTGCCCATGTGA |
| Rev primer 5' integration *Ckm*-/*Mb*-GFP | GTCTCGTGGGCTCGGAGATGTGTATAAGAGACAGATCAGCGCAGAAGTCTCAG |
| Rev primer 3' integration *Ckm*-GFP | GTCTCGTGGGCTCGGAGATGTGTATAAGAGACAGGGTTCTCCCCCTGTGAACCT |
| Rev primer 3' integration *Mb*-GFP | GTCTCGTGGGCTCGGAGATGTGTATAAGAGACAGGCCCCATGGTGGCTTCTAAA |
| Fwd primer 3' integration *Ckm*-/*Mb*-GFP | TCGTCGGCAGCGTCAGATGTGTATAAGAGACAGTCTTAGTCAGGCTGCCCCTC |
| Rev primer cDNA *Ckm*-/*Mb*-GFP | GTCTCGTGGGCTCGGAGATGTGTATAAGAGACAGGGACTCACACTGATCTCCATC |
| Fwd primer cDNA *Ckm*-GFP | TCGTCGGCAGCGTCAGATGTGTATAAGAGACAGACCTCCACAGCACAGACAGAC |
| Fwd primer cDNA *Mb*-GFP | TCGTCGGCAGCGTCAGATGTGTATAAGAGACAGCATCCTTGTCCCTGTGGGTG |
| **qPCR primers** | |
| Fwd *hF9* | GATAGAGCCTCCACAGAATGC |
| Rev *hF9* | TTGGATAACATCACTCAAAGCAC |
| Probe *hF9* | FAM/TCCCTTGGC/ZEN/AGGTTGTTTTGAATGG |
| Fwd *Hprt* | CTGGTGAAAAGGACCTCTCGAAG |
| Rev *Hprt* | CCAGTTTCACTAATGACACAAACG |
| Fwd *Actb* | GGCTGTATTCCCCTCCATCG |
| Rev *Actb* | CCAGTTGGTAACAATGCCATGT |
| Fwd *Ppia* | GGGTGGTGACTTTACACGCC |
| Rev *Ppia* | CTTGCCATCCAGCCATTCAG |
| Fwd *Gapdh* | CCTCGTCCCGTAGACAAAATG |
| Rev *Gapdh* | TGAAGGGGTCGTTGATGGC |
| **UDiTas** | |
| GSP *Ckm* intron | GTCTCGTGGGCTCGGAGATGTGTATAAGAGACAGCAGCCCCAGATCCTGTATTTTTG |
| GSP GFP insert | GTCTCGTGGGCTCGGAGATGTGTATAAGAGACAGATCAGCGCAGAAGTCTCAG |
| i5 | AATGATACGGCGACCACCGAGATCTACACTCGTCGGCAGCGTC |
| **5'RACE** | |
| GSP RT primer *hF9* | CTCCTTCATGGAAGCCAGCA |
| GSP PCR primer *hF9* | TCATTGCACACTCTTCACCC |
| TSO | GCTAATCATTGCAAGCAGTGGTATCAACGCAGAGTACATRGRGRG |
| TSO-specific primer | CATTGCAAGCAGTGGTATCAAC |
